# Direct Reciprocity Under Uncertainty Does Not Explain One-Shot Cooperation, But It Can Explain Norm Psychology

**DOI:** 10.1101/001446

**Authors:** Matthew R. Zefferman

**Affiliations:** Graduate Group in Ecology, University of California, Davis National Institute for Mathematical and Biological Synthesis, University of Tennessee, Knoxville

## Abstract

Humans in many societies cooperate in economic experiments at much higher levels than would be expected if their goal was maximizing economic returns even when interactions are anonymous and one-shot. This is a puzzle because paying a cost to benefit another player in one-shot interactions has no direct benefit to the cooperator. This paper explores the logic of two competing evolutionary hypotheses to explain this behavior. The “norm psychology” hypothesis holds that a player’s choice of strategy is heavily influenced by socially-learned cultural norms. Its premise is that over the course of human evolutionary history, cultural norms varied considerably across human societies and through a process of gene-culture co-evolution, humans evolved mechanisms to learn and adopt the norms of their particular society. The “evolutionary mismatch” hypothesis holds that pro-social preferences evolved genetically in our hunter-gatherer past where one-shot anonymous interactions were rare and these evolved “protocols” for cooperation are misapplied in modern, laboratory, conditions. I compare these hypotheses by adopting a well-known model of the mismatch hypothesis. I show that the cooperation generated by the model is based on a flawed assumption - that the best thing to do is cooperate in a repeated game. I show that repeated games generate a great diversity of behavioral equilibria, in support of the norm psychology hypothesis’s premise. When interaction is repeated, adopting local norms is a more evolutionarily successful strategy than automatically cooperating. If various groups are at different behavioral equilibria, then cultural selection between groups tends to select for cooperative behavior.

## 1. Introduction

A puzzling finding in experimental economics is that in many societies humans cooperate in laboratory experiments at much higher levels than they would if they were money-maximizing agents. Players act as though they have “pro-social preferences,” that is, in addition to their own welfare, they care about some combination of the welfare of other players, fairness and equality (Fehr and Schmidt, 1999; Bolton and Ockenfels, 2000). The strongest evidence for pro-social preferences come from simple games, such as the “dictator game,” where participants are given an amount of money that they can divide between themselves and an anonymous stranger. While the money-maximizing agent would keep the entire sum, players consistently distribute substantial sums to anonymous strangers (Camerer, 2003). Pro-social play have also been observed in more complicated games, such as ultimatum games, trust games, Prisoner’s Dilemmas, and public goods games (Fehr and Schmidt, 1999; Bolton and Ockenfels, 2000; Camerer, 2003) and has been documented in many societies, though there there is substantial variation both within and between societies (Henrich et al., 2005). Pro-social play is especially puzzling when a game is played only once with an anonymous partner since one-shot anonymous interactions eliminate reciprocity and reputation-building as motivations. What explains the existence of pro-social play in one-shot anonymous games?

In this paper I explore two competing evolutionary hypotheses for the origins of pro-social play in one-shot economic experiments. One, the “norm psychology hypothesis,” is that cooperative play is primarily due to pro-social norms acquired during a human’s life through social learning (Richerson and Boyd, 2005; Boyd and Richerson, 2009; Chudek and Henrich, 2011). This hypothesis is premised on the proposition that over the course of humans’ evolutionary history, the social norms of different human groups were *highly varied and subject to frequent change*. Therefore, it would have been difficult for genetic adaptations in response to *specific* norms to take hold. However, humans with a “norm psychology” that helped them better learn and conform to the prevailing norms of their *particular* society would do better than those without this ability. Variation in social norms was partly due to local geographical and ecological circumstances, but also to direct reciprocity, altruistic punishment (strong reciprocity) and reputation-building (indirect reciprocity) which can generate and reinforce a very wide range of equilibria (Boyd and Richerson, 1992; Boyd, 2006). This hypothesis is premised on the condition that different groups of individuals engaged in repeated interaction will evolve to *many different* behavioral equilibria, some of which will be more cooperative than others.

Another hypothesis, often called the “mismatch hypothesis,”^1^ is that co-operative play is primarily due to genetic programs, mechanisms or protocols acquired in ancient times through selection on genes (Kanazawa, 2004; Hagen and Hammerstein, 2006; Price, 2008; Delton et al., 2011; Pinker, 2012; Krasnow et al., 2012; McCullough et al., 2013; Pedersen et al., 2013). This hypothesis is premised on the proposition that over the course of humans’ evolutionary history, there were very few one-shot or anonymous encounters and repeated interactions created to *consistent and persistent cooperation*. Humans who were endowed with a propensity to cooperate in one-shot interactions would do better because they would more easily capture the benefits of future interactions that would undoubtedly follow. This hypothesis predicts that groups of individuals engaged in repeated interaction will consistently evolve to cooperative equilibria. In laboratory experiments, cooperative play from an “evolutionary mismatch” between these cooperative genetic protocols and modern experimental settings. In other words, cooperative play in one-shot economic games results from our “modern skulls housing a stone age mind” (Cosmides and Tooby, 1997).

Table 1 summarizes key differences between the norm psychology and mismatch hypotheses. Fehr and Henrich (2003) and Henrich et al. (2004) have raised empirical objections to the mismatch hypothesis based primarily on ethnographic and experimental evidence. However, others have found these objections unconvincing (Hagen and Hammerstein, 2006) and question the plausibility of the norm psychology hypothesis as an alternative (Price, 2008). This empirical debate is beyond the scope of this paper. Instead, I focus on which hypothesis is better supported by theory. Towards this end, I adopt a model developed to formally explain and support the mismatch hypothesis (Delton et al., 2011), showing that if certain artificial constraints on the scope of behaviors available to selection are relaxed, it provides better support for the norm psychology hypothesis.

**Table 1:**
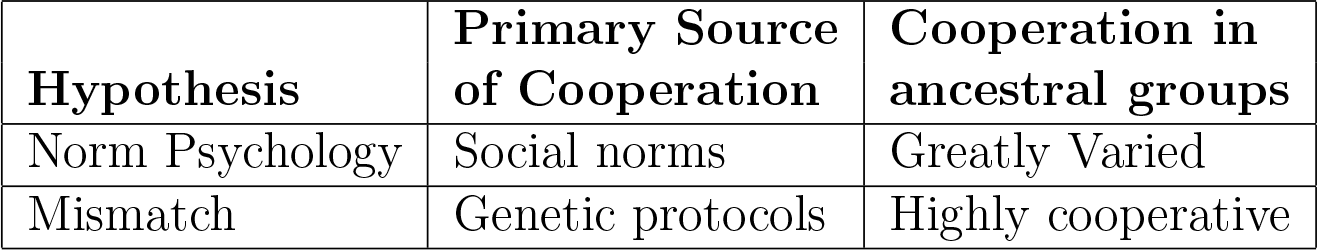
Key differences between the norm psychology and mismatch hypotheses

### 1.1. The DKCT Model

In this paper, I adopt a recent model by Delton, Krasnow, Cosmides and Tooby (2011) (hereafter “DKCT” and “the DKCT model”) which has sought to put the mismatch hypothesis on more solid theoretical footing. Like other proponents of the mismatch hypothesis, DKCT hypothesize that one-shot cooperation can be explained by genetic selection among “our band-living hunter-gatherer ancestors” (Delton et al., 2011, S1) whose lives were dominated by repeated interactions. However, they add a twist. When strangers meet there is *uncertainty* about whether it will be a one-time encounter or a repeated interaction. Since repeated interaction can create, over time, greater absolute costs and benefits than one-shot interactions, mistaking a repeated interaction for a one-shot interaction is more costly than mistaking a one-shot interaction for a repeated one (see also Krasnow et al. (2012), Pinker (2012), McCullough et al. (2013), and Pedersen et al. (2013)). They run a series of simulations of this idea and find that, as premised by the mismatch hypothesis, agents will evolve a propensity to cooperate, even if there is strong evidence that an interaction is one-shot. In this section I briefly describe the DCKT model, which is further elaborated in the supplemental materials of their paper (Delton et al., 2011). In the subsequent sections, I describe reasons one might be skeptical of their results based on previous theory and show how the high levels of one-shot cooperation they find are an artifact of constraining their agents to an evolutionary history that allows only two of an infinite number of possible strategies. Then I show how groups exposed to different evolutionary histories in the same model will have highly varied patterns of behavior, as premised by the norm psychology hypothesis. Wherever I was unsure of the original model’s details, I consulted the authors who helpfully provided clarification.

In the DKCT model, 500 agents are born, randomly pair into dyads, play a game, reproduce, and die. They play either a one-shot or repeated Prisoner’s Dilemma, where, in each round, they can pay a cost, *c*, to confer a benefit, *b*, to their partner (Figure 1). With a probability, *P*, the game is one-shot. Otherwise it is repeated. If the game is repeated, after each round the probability that the game lasts another round is *w*. Therefore, given a repeated game, the number of rounds is drawn from a geometric distribution with an expectation of 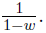

**Figure 1:**
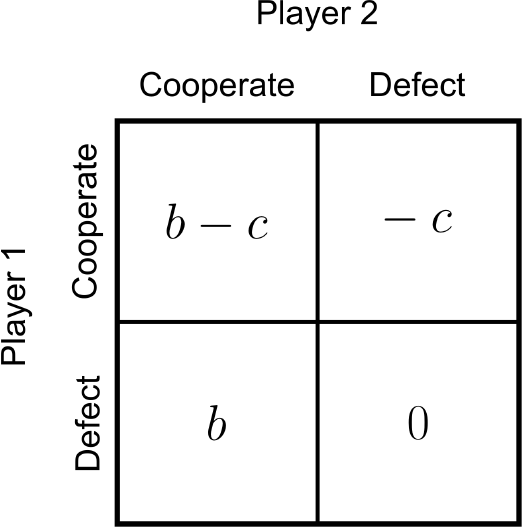
The payoffs for Player 1 in one round of a repeated Prisoner’s Dilemma game. In this game an agent who cooperates pays a cost *c* to provide a benefit *b* to its partner. An agent who defects does not. Since defecting always gives a higher payoff than cooperating, agents should always defect in a one-shot game if their goal is to maximize returns. In a repeated game, the same pair of agents play this game multiple times.

Before playing the game each agent in a dyad independently receives an imperfect signal about whether the interaction is likely to be repeated or one-shot. As shown in Figure 2, if the game is repeated, the signal is drawn from a normal distribution with a mean of *d/*2 and a standard deviation of one. If the game is one-shot, the signal is drawn from a normal distribution with a mean of −*d/*2 and a standard deviation of one. Since these distributions overlap, an agent cannot be sure which distribution the signal was drawn from. Since the amount of overlap decreases with the size of *d*, this parameter is a measure of the certainty in the model.

**Figure 2:**
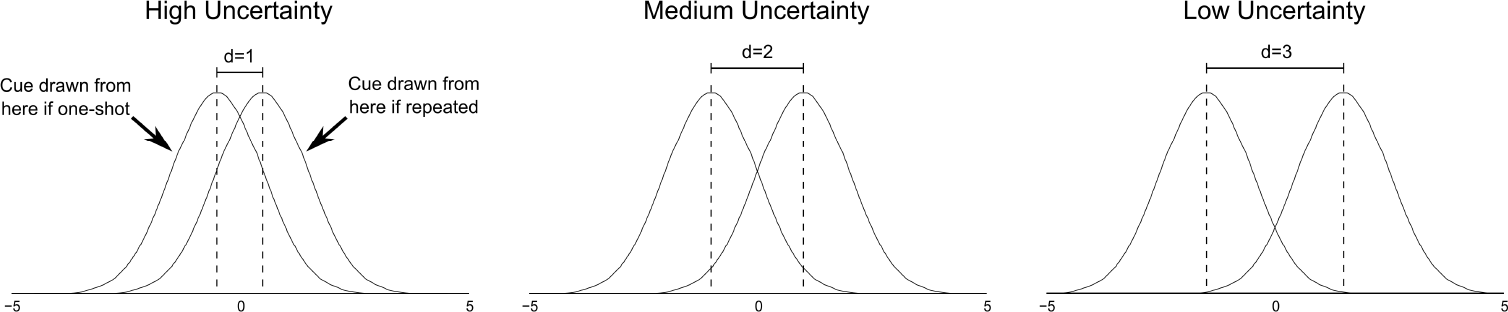
Nature draws a signal from one of two distributions depending on whether an interaction is one-shot or repeated. If a game is repeated, the signal is drawn from a normal distribution with a mean of *d/*2 and a standard deviation of one. Otherwise the signal is drawn from a distribution with a mean of −*d/*2. DKCT use three values of *d* with lower values indicating more uncertainty since there is more overlap between distributions.

Each agent is born with a “cue threshold,” which is a number it uses to pick a strategy based on the imperfect signal it receives. If an agent’s signal is greater than its cue threshold (indicating a repeated game), the agent plays Tit-for-Tat (TFT), a strategy that cooperates on the first round of play and thereafter repeats the actions of the other agent on the previous round. If an agent’s signal is less than its cue threshold (indicating a one-shot game), the agent plays Always Defect (ALLD), a strategy that defects on every round.

In the first generation, cue thresholds are distributed normally with a mean of 0 and a standard deviation 0.025. When an agent is born in later generations, it inherits a cue threshold from a member of the previous generation, with a probability proportional to the members’ relative payoffs. However there is a high, 5%, chance that an agent’s decision threshold will mutate, changing by a normally distributed random variable with a mean of 0 and a standard deviation 0.025. After reproducing, all members of the previous generation are removed from the population. DKCT’s simulations each lasted for 10,000 generations. Agents start each generation with a baseline payoff of 10.

DKCT run their simulations under 750 separate parameter conditions which are given in Table 2. They found that agents evolved high frequencies of one-shot cooperation for many of these 750 parameter combinations. In fact, they often developed a strong bias towards playing TFT even when there was strong evidence that an interaction is one-shot.^2^ However, the agents in DKCT’s simulations can only play two strategies, TFT and ALLD and, as I described below, a more complete set of possible strategies should generate a diversity of behavioral equilibria. I then show that if other strategies can invade TFT in DKCT’s model, one-shot cooperation can virtually disappear from the population, demonstrating the flaw in the assumption that repeat interaction leads to cooperative equilibria.

**Table 2:** Table of parameters. DKCT and I both run our simulations under every combination of these parameters, for 750 total simulations. In addition, I run the model where each agent has only one partner in their lifetime (as in DKCT) and where each agent has ten lifetime partners.

| Symbol | Definition | Values |
| --- | --- | --- |
| $c$ | Cost of cooperating in a round | 1 |
| $b$ | Benefits from cooperation to other player in one round | 1.5, 2, 3, 4, 5, 6, 7, 8, 9, 10 |
| $P$ | Probability that an interaction is one-shot | 0.1, 0.3, 0.5, 0.7, 0.9 |
| $w$ | In a repeated interaction, probability of a subsequent round | 0.5, 0.7, 0.9, 0.95, 0.99 |
| $d$ | Amount of uncertainty over whether a game is one-shot or repeated | 1, 2, 3 |

### 1.2. Repeated Games Create Many Unstable Equilibria

Repeated games do not necessarily favor cooperation, but have many behavioral equilibria, some cooperative and others not. DKCT chose TFT for their simulations in part because it “has the additional benefit of being familiar to most readers.” TFT is well-known because it famously performed better than any other strategy submitted to two computerized tournaments run by Robert Axelrod (Axelrod and Hamilton, 1981; Axelrod, 1984). TFT was one of the simplest strategies entered and since it was both “nice” (cooperating on the first turn) and “retaliatory” (defecting after a partner defects), its success seemed to cemented “niceness” and “retaliation” as the paths to evolutionary success in repeated games.

However, TFT’s success hinged on the particular mix of strategies entered in the tournaments and it is not generalizable to other mixes of strategies. One reason is explained by the “folk theorem” of repeated games^3^ which shows that if the Prisoner’s dilemma is sufficiently repeated, *any* pattern of behavior can be an equilibrium (Rubinstein, 1979; Fudenberg and Maskin, 1986). All that is required is that players adhere to a pattern and withhold cooperation from any other player that deviates from the pattern. This is a problem for the premise that repeated interactions enviably lead to cooperative equilibria. It is not enough to assume that cooperation is a possible outcome of a repeated interaction, as both cooperative and non-cooperative equilibria are possible. As Boyd (2006) puts it, “when everything is an equilibrium, showing that reciprocity is an equilibrium too does not really tell you much.”

Another reason why cooperation is not the inevitable outcome of repeated Prisoner’s Dilemma is that any equilibrium is unstable, that is it can be invaded by other strategies. This was quickly pointed out in the case of TFT (Williams, 1984; Selton and Hammerstein, 1984), after Axelrod and Hamilton (1981)’s initial claims of its stability. It was later proved that there are *no* evolutionarily stable strategies in a sufficiently repeated Prisoner’s Dilemma (Boyd and Lorberbaum, 1987; Farrell and Ware, 1989; Lorberbaum, 1994). For example, Boyd and Lorberbaum (1987) show that TFT can be invaded by a more forgiving strategy called “Tit-for-Two Tats” (TF2T) that is nice but only retaliates after its opponent defects twice in a row, in combination with a nasty strategy called “Suspicious Tit-for-Tat”(STFT) that is similar to TFT except that it defects on the first turn. TF2T increases in the population relative to TFT since it has higher payoffs when playing STFT. However, STFT exploits TF2T’s forgiveness and eventually invades to fixation. This is an example of selection in repeated interactions destabilizing a nice strategy, like TFT, and bringing a “nasty” strategy, STFT to fixation. DKCT do not allow for this type of invasion in their simulations, which increases the amount of cooperation (TFT) in their model.

This is backed up by computational modeling. For example, when Axelrod (1997, 21-22) used genetic algorithms to evolve strategies to play against a representative sample of the strategies submitted to his tournament, sometimes strategies similar to TFT would evolve, but sometimes strategies that performed better than TFT evolved, and all of these strategies defected on the first move. Similarly,(Nowak and Sigmund, 1989) showed that, even in a set of fairly very simple strategies, the evolutionary dynamics are complicated and would cycle between “nice” and “nasty” strategies. In the next sections, I show that DKCT’s inclusion of uncertainty does not make their model to invasion by non-cooperative strategies. This suggests that evolving a strong bias towards cooperating on the first interaction with a stranger is unlikely to be beneficial in the long term.

## 2. Methods

DKCT’s simulations all start with a very specific behavioral proposition. Agents will always defect if given sufficient evidence that an interaction is one-shot and play TFT if given sufficient evidence than an interaction will be repeated. However, as explained above, theory indicates that any behavior is a plausible equilibrium in a repeated PD. DKCT’s model is different than a standard repeated PD because they include a degree of uncertainty over whether a game is one-shot or repeated. Does this change the logic of repeated games?

To answer this question, I replicated the DKCT model with different combinations of initial strategies. To ensure that any differences between outcomes of these combinations were due only to the mix of strategies, I also replicated the aspects of DKCT’s model that were particularly friendly to generating cooperative outcomes. First, all interactions involve only two players, which is the condition where reciprocity most easily generates cooperation. Reciprocity because rapidly less effective as more players are added to a game (Joshi, 1987; Boyd and Richerson, 1988). Second, the population *begins* with every agent playing a cooperative strategy when there is enough evidence that a game is repeated. This is a strong assumption because the more likely ancestral condition would have little to no altruistic cooperation and it is much harder for reciprocity to explain cooperations *origins* than its maintenance. Third, the parameter values were very friendly to cooperation. A single cooperative act could have up to a 1000% return on investment and this high return could potentially be realized over hundreds of interactions. However, even keeping all of these cooperation-favoring assumptions, I find that the amount of one-shot cooperation varies widely.

I ran each simulation under two conditions for the number of partners an agent has in its lifetime. In one condition, as in DKCT’s original simulations, each agent only interacts with one other agent in its lifetime. In the other condition, I increase the number of dyadic partners an agent has to ten. This is not only more realistic, since most humans interact with multiple other people in their lifetime, but it also decreases the variance in payoffs due to the number of rounds each agent plays. A quirk of having only one partner in these simulations is that under some parameter conditions (i.e., high *P* and *w*) the number of rounds agents play is highly skewed where almost every dyad plays only one round, but a very small subset of dyads may play in the hundreds of rounds. This creates stochastic shocks where low-performing strategies jump to near fixation in a generation simply because a dyad was randomly assigned a game with substantially more rounds than all other dyads combined. However, as agents interact with more dyadic partners, the highly skewed distribution of numbers of rounds played smooths out due to the central limit theorem (see Appendix C). I present the results of both conditions in Appendix A.

I ran simulations of the DKCT model for all 750 of their parameters, and for both the one-partner and ten-partner cases, under three initial conditions. To demonstrate consistency with DKCT’s original findings, the first condition is an exact replication of the DKCT’s simulations which include only TFT and ALLD. In the other two conditions, I introduce strategies not considered by DKCT. Since written descriptions of strategies for repeated games can sometimes be ambiguous, I precisely represent the strategies of all treatments, following Rubinstein (1986) and Miller (1996), as Finite State Automata in Appendix D.

### 2.1. Treatment 1: TFT and ALLD

In Treatment 1, as in DKCT’s model, agents play TFT if the cue is above their cue threshold (indicating a higher probability of a repeated game) and ALLD if the cue is below their threshold (indicating a higher probability of a one-shot game). Figure 3A shows the expected payoffs for one-shot and repeated games for agents playing TFT and ALLD. I replicate their model for all 750 combinations of parameter values explored by DKCT. I replicated this treatment with agents having only one partner (as conducted by DKCT) and with each agent having ten partners.

**Figure 3:**
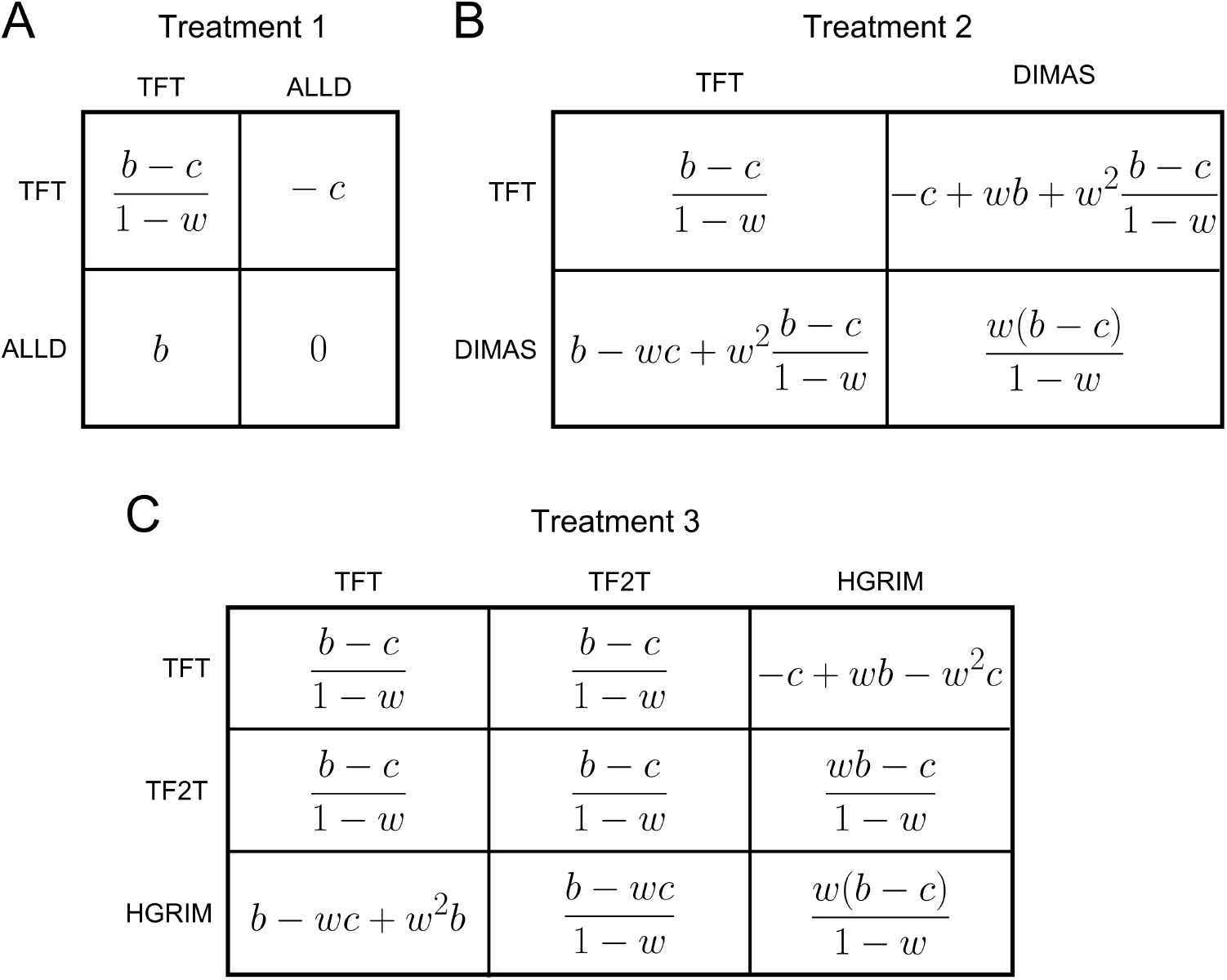
Expected payoffs for each strategy in the three treatments, assuming a repeated game. *b* is the benefits to cooperation, *c* is the cost of cooperating, and *w* is the probability that, after each round in a repeated game, there is another round. Note that the number of rounds in a repeated game is geometrically distributed where the expectation of the number of rounds is 1*/*(1 − *w*). **A** shows the expected payoffs for Treatment 1. **B** shows the expected payoffs for Treatment 2. **C** shows the expected payoffs for Treatment 3.

### 2.2. Treatment Two: TFT and DIMAS

In DKCT’s original simulations first-round defectors must continue, by assumption, to defect for all time. In other words, agents cannot repent. However, the ability to make amends is an important part of most humans’ behavioral repertoires. McNally and Tanner (2011) suggest that a repentant strategy would perform better than ALLD in DKCT’s model, reducing the amount of one-shot cooperation. To test this suggestion, I replace ALLD with a simple strategy that defect on the first round, but finding itself in a repeated game, immediately repents and begins cooperating. I dub this strategy “DIMAS” after the biblical thief whose repentance earns him eternal rewards in paradise.

Figure 3B shows the expected payoffs for one-shot and repeated games for agents playing TFT when the signal is above their cue threshold and DIMAS when the signal is below it. These are similar to those in Figure 3A except that DIMAS typically earns higher payoffs than ALLD because it both cooperates with itself starting in round two and cooperates with TFT starting in round three.

### 2.3. Treatment Three: TFT, TF2T and HGRIM

In the third treatment I introduce a more savvy repentant strategy and a more forgiving cooperative strategies, reflecting previous work (Boyd and Lorberbaum, 1987) on forgiveness and repentance.^4^ I replace ALLD with a savvy repentant strategy dubbed “Hesitant Grim” (HGRIM). HGRIM defects on the first round, cooperates on the second, and then plays a trigger strategy where it cooperate until its opponent defects, and then continues to defect thereafter. HGRIM is still a fairly simple strategy, comparable in complexity to TF2T (both can be represented as three-state Finite State Automata as shown in Appendix D). HGRIM differs from DIMAS in that, while it still cooperates after the first round with repentant and forgiving strategies, it does not cooperate with unrepentant or retaliatory strategies.

To simulate the invasion of a novel cooperative strategy, I replace a small fraction (5%) of the initial population with agents who play a slightly more forgiving strategy Tit-for-Two-Tats (TF2T) where others would play TFT. TF2T is often a high-performing strategy. In fact, Axelrod submitted it himself to his second tournament after determining that it would have won the first, had it been entered (Axelrod, 1984).^5^

Having two nice strategies means that selection now acts on two traits in the model. The first is the cue threshold, as in the other treatments, and the second is the nice strategy (TFT or TF2T) employed with sufficient evidence of a repeated interaction. Therefore, in Treatment 3 each agent has two separate parents from the previous generation (instead of one parent as in Treatments 1 and 2). Each parent is chosen with a probability proportional to their relative payoff. Each agent inherits a cue threshold and a nice strategy from one of its parents drawn randomly and independently for each trait. There is also a 0.1% chance that an agent’s nice strategy will mutate from TFT to TF2T or *vice versa*.

## 3. Results

The three treatments described in Section 2 represent the DKCT model under slightly different mixes of starting strategies. Biasing the results towards DKCT’s findings, in all three treatments agents play nasty strategies with enough evidence that an interaction is one-shot and nice strategies with enough evidence that an interaction is repeated. The nice strategy played by agents in the first two treatments is TFT, as in DKCT’s simulations, and in the third treatment 95% of agents initially play TFT. Despite these similarities in initial conditions, the expected frequency of one-shot cooperation varies widely between treatments, with one-shot cooperation fairly frequent under many parameter combinations in Treatment 1 and almost entirely absent in Treatment 3.

Figure 4 illustrates how levels of one-shot cooperation vary dramatically when different strategies are in the population. Figure 4A shows that when agents are constrained to ALLD and TFT, as in DKCT’s simulations, relatively high levels of one-shot cooperation can evolve. These results are similar to those in Delton et al. (2011)’s original simulations which is unsurprising, since their strategy space was constrained to the same two strategies. As shown in Appendix Appendix A, this result holds for most of DKCT’s 750 parameter combinations. If an agent born into a society with an evolutionary history similar to Treatment 1 for these parameter combinations, it would do better if it learned and employed a strategy of one-shot cooperation. But is this true for groups with other evolutionary histories?

**Figure 4:**
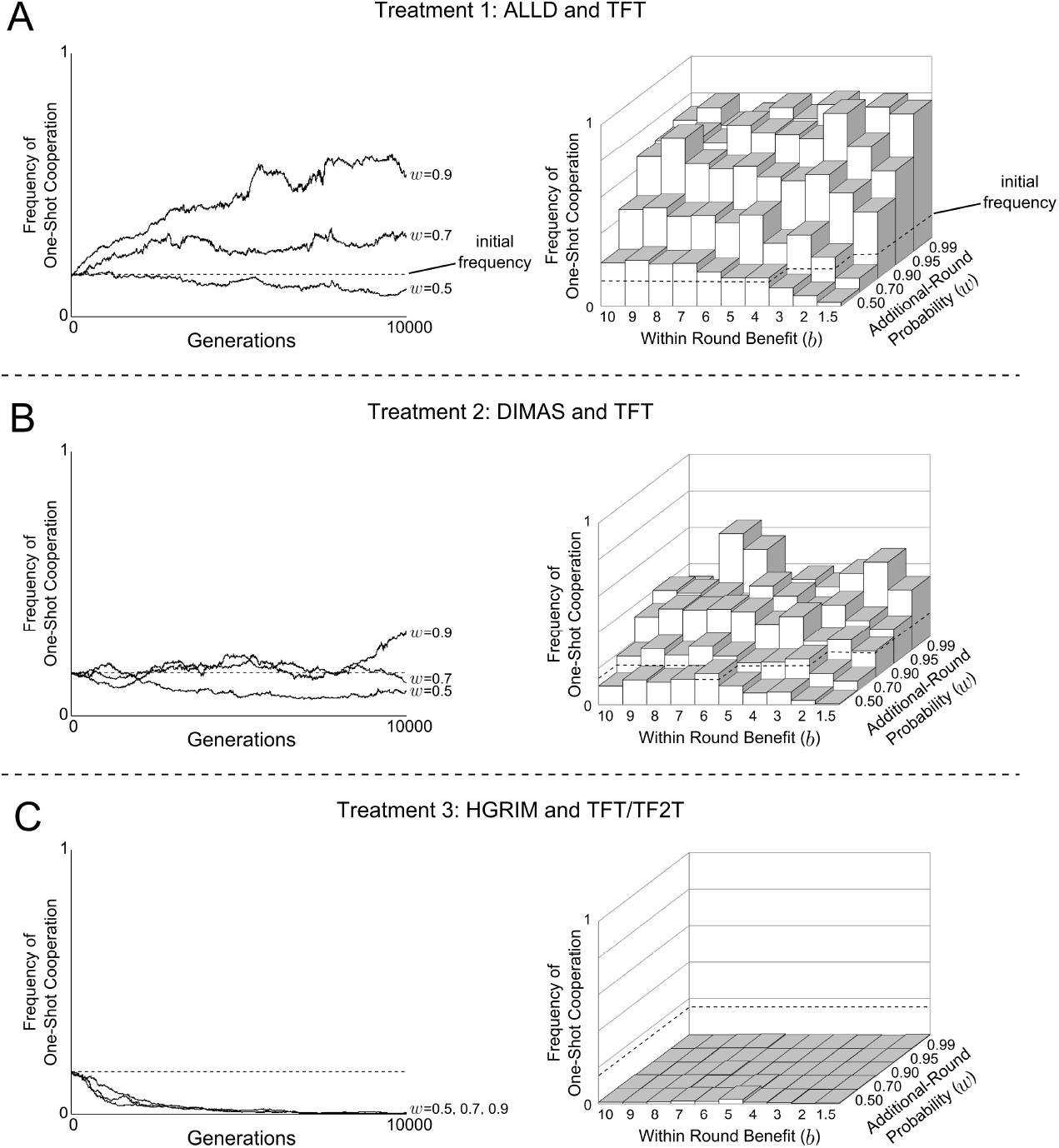
The expected frequency of one-shot cooperation substantially decreases with the addition of forgiving and repentant strategies. This figure shows both time-series and final expected frequencies of one-shot cooperation for selected parameter combinations (*P* = 0.5, *d* = 2) for Treatment 1 (A), Treatment 2 (B) and Treatment 3 (C). (These parameter combinations match Delton et al. (2011)’s Figure 3.) Although A shows high expected frequencies of one-shot cooperation as reported by DKCT, C shows that this virtually disappears when repentant and forgiving strategies invade. B shows that a forgiving strategy alone is somewhere in-between. This suggests that, not only will there be different expectations of cooperation in different societies, but that DKCT’s simulations were at the higher end of the cooperative scale. This general pattern holds across the 750 parameter combinations as shown in Appendix A.

In Treatment 3, as shown Figure 4B and Appendix A, there is a dramatically different result. When agents employ a savvy repentant strategy, HGRIM, and a forgiving strategy, TF2T, is able to invade the population, one-shot cooperation based on TFT is dislodged under all parameter combinations with one-shot defection eventually taking over the system. In fact, agents evolve to defect on the first round even when there is strong evidence that an interaction is repeated. An agent born into a society with an evolutionary history similar to Treatment 3 would do best if it employed a strategy of one-shot defection. This is the opposite strategy of what an agent would best employ in Treatment 1. The results of Treatment 2, as shown Figure 4C and Appendix A, are somewhere in-between those of Treatment 1 and Treatment 3.

This general pattern holds for all 750 parameter combinations as shown in Appendix A. One-shot cooperation is generally common in groups with evolutionary histories similar to Treatment 1, uncommon in groups with histories similar to Treatment 3 and intermediate in groups with histories similar to Treatment 2. Interestingly, even though initially the number of agents playing TFT is very high, for all parameter conditions one-shot cooperation decreases well below initial conditions for Treatment 3 and, in most cases, virtually disappears. This effect is clear when, as below, the case of maximum uncertainty is examined analytically.

## 4. The Case of Complete Uncertainty

DKCT credit the high expected frequency of one-shot cooperation in their simulations to the *uncertainty* over whether an interaction is one-shot or repeated. The strongest case for one-shot cooperation in their model is when uncertainty is the greatest that is when players have no indication as to whether they are in a one-shot or repeated game, that is, there is no signal. This case reduces to a standard repeated Prisoner’s Dilemma where the probability of transitioning from the first to second round is (1 − *P)w*, which is lower than the transition probability for every other round, *w*. In this section I give the conditions where one-shot cooperation is stable and the conditions where it is risk-dominant when the strategies available to selection mirror the three treatments described above. However, these results only apply to the case where no novel strategies invade. As described above, any stable equilibrium described here could be invaded by the right mix of novel strategies.

Table 3 shows that the conditions favoring one-shot cooperation decrease when there are repentant and forgiving strategies. This can been seen by counting how many of the 250 combinations of *b*, *w* and *P* that DKCT specify as plausible in their paper stabilize one-shot cooperation and in how many combinations one-shot cooperation is risk-dominant. Stability is required for risk-dominance and when there are more than one stable equilibria, a population will, in the long run, spend more time at the risk-dominant one.^6^ For example when selection only acts on TFT and ALLD (as in DKCT’s model and my Treatment 1), TFT is stable under a high number, 85%, of the plausible parameter combinations and is also risk-dominant in a majority 65% of them. However when ALLD is replaced with DIMAS, TFT is stable in only 64% of the plausible parameter combinations and is risk-dominant in only 42% of them. This suggests that the frequency of one-shot cooperation in Treatment 2 is more dependent on environmental factors (parameter combinations) than in Treatment 1.

**Table 3:**
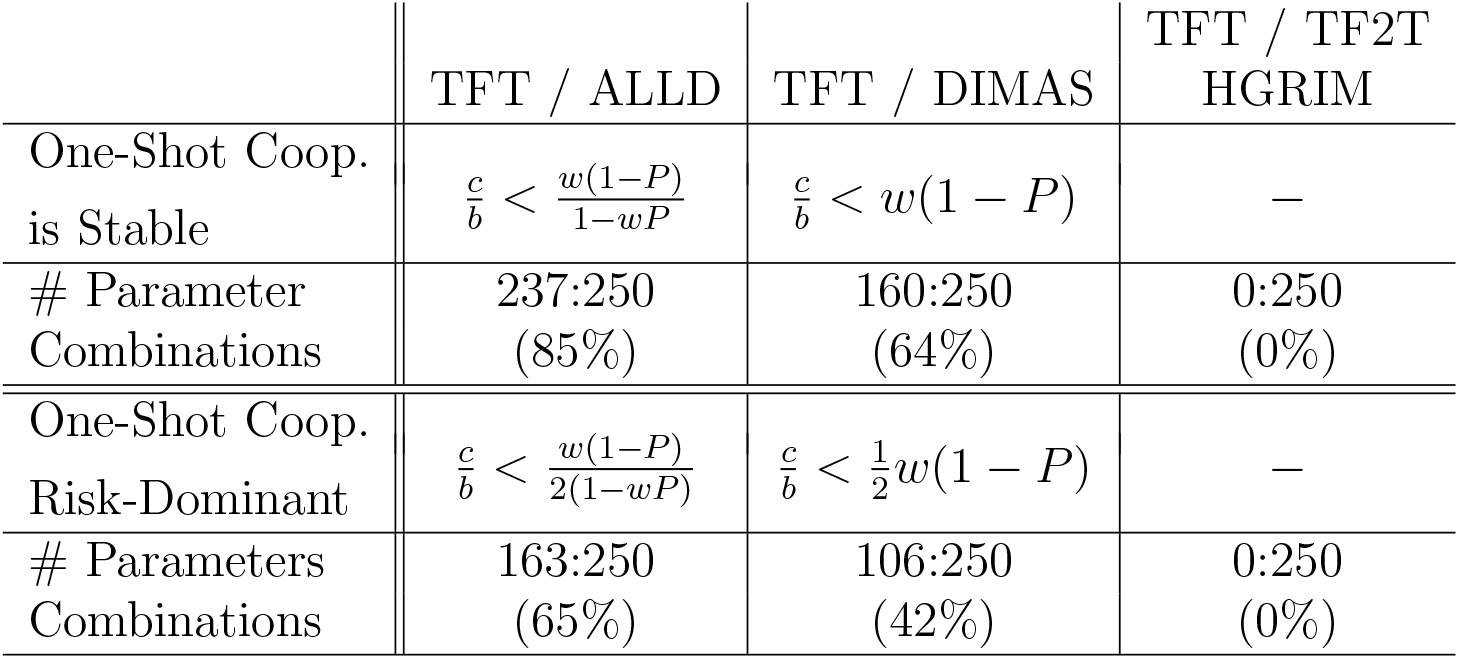
The number of parameter conditions where one-shot cooperation is stable and risk dominant decrease if repentant and forgiving strategies are allowed in the population. When a repentant and forgiving strategy can invade a population with TFT, as in Treatment 3, one-shot cooperation goes to zero.

|  | TFT / ALLD | TFT / DIMAS | TFT / TF2T<br>HGRIM |
| --- | --- | --- | --- |
| One-Shot Coop.<br>is Stable | $\frac{c}{b} < \frac{w(1-P)}{1-wP}$ | $\frac{c}{b} < w(1-P)$ | — |
| # Parameter<br>Combinations | 237:250<br>(85%) | 160:250<br>(64%) | 0:250<br>(0%) |
| One-Shot Coop.<br>Risk-Dominant | $\frac{c}{b} < \frac{w(1-P)}{2(1-wP)}$ | $\frac{c}{b} < \frac{1}{2}w(1-P)$ | — |
| # Parameters<br>Combinations | 163:250<br>(65%) | 106:250<br>(42%) | 0:250<br>(0%) |

Finally, in a population with HGRIM and TF2T (as in Treatment 3 above) one-shot cooperation is *never* stable, even in an uncertain environment. And since risk-dominance requires stability, TFT is never risk-dominant. We would expect that there would be no one-shot cooperation in such a population (a finding reflected in my discussion of Treatment 3 in Section 3). As I show in Appendix B, this is not only the case for the DKCT’s plausible parameter conditions, but for *all* possible values of *b*, *w*, and *P*. That one-shot cooperation is so readily replaced by one-shot defection when a population includes repentant and forgiving strategies, even under conditions of complete uncertainty, is a problem for the mismatch hypothesis which is premised on reciprocity leading inevitably to cooperative equilibria. Instead, it further supports the premise of the norm psychology hypothesis that there will be high behavioral variation between groups that critically depend on a particular society’s evolutionary history.

## 5. The Evolution of Norm Psychology

I have demonstrated that repeated interaction, even under uncertainty, does not necessarily favor one-shot cooperation. Instead, it supports a variety of path-dependent equilibria, depending on the evolutionary history of the population, which are the conditions that favor a norm psychology. For example, I explored three different evolutionary trajectories. In all of these trajectories, the “nasty” strategy is evolutionarily stable given the mix of other strategies in the population and in some, both the nice and the nasty strategies are evolutionarily stable. In Treatment 1, if ALLD is at fixation in a population, playing ALLD has a higher payoff than playing TFT. Similarly, in Treatments 2 or 3, when DIMAS or HGRIM are at fixation, playing DIMAS or HGRIM has higher payoff than playing TFT, ALLC or TF2T. When TFT is an evolutionarily stable strategy (85% of parameter combinations in Treatment 1, 65% in Treatment 2 and 0% in Treatment 3) playing TFT has a higher payoff than the nasty strategy. In all cases, when one’s group is at equilibrium the same or higher payoff is achieved by adopting the equilibrium strategy than having predisposition to play a cooperative strategy. Since groups can undergo a large number of evolutionary trajectories, individuals born into a group will be better off adopting the norms of the group, including first round cooperation or defection, than they would automatically playing a strategy like TFT.

## 6. Discussion

This paper compares the logical underpinnings of two competing hypotheses seeking to explain the prevalence of cooperation in one-shot laboratory experiments. The mismatch hypothesis, which is based on the premise that repeated interactions inevitably lead to one-shot cooperation, found support in a recent model. However, the high frequencies of one-shot cooperation that evolved in the model were due to agents’ evolutionary possibilities constrained to TFT and ALLD. When agents can employ repentant and forgiving strategies in the same model, the frequency of one-shot cooperation falls and, in some scenarios, disappears for all plausible parameter conditions and degrees of uncertainty. This suggests that societies undergoing independent evolutionary trajectories will have very different levels of one-shot cooperation. This sets the conditions for the norm psychology hypothesis which is premised on the idea that humans evolved, through a process of gene-culture coevolution, to flexibly learn and adopt the particular norms of their particular society.

The norm psychology hypothesis is also supported by the wide variation in levels of cooperative play in economic experiments observed across societies (Henrich et al., 2004, 2005). In fact, two society-level measurements, a society’s degree of market integration and scale of potential payoffs to cooperation outside of the lab, are much better predictors of cooperative play in economic experiments than ndividual-level measures (Henrich et al., 2005). In addition, the mismatch hypothesis does not explain human cooperation that is is non-dyadic collective action since direct reciprocity is not effective at producing altruistic cooperation equilibria in most *n*-person interactions where *n* is greater than around two (Joshi, 1987; Boyd and Richerson, 1988).

How does the norm psychology hypothesis explain the large amount of cooperation in many societies? The norm psychology hypothesis has a close cousin, the “cultural group selection hypothesis” which focuses less on the origins of learning mechanisms in individual human behavior than on the population-level consequences of these mechanisms.

The hypothesis, first proposed by Charles Darwin (1873), is that when groups of humans are in competition, groups where individuals are more cooperative with other group members will tend to out-compete groups where cooperation is rare. For example, consider groups undergoing cultural evolution under complete uncertainty in a strategy space similar to that in Treatment 2 - where agents either play TFT or DIMAS. Consider a situation where each of these strategies can be a stable equilibria and neither is risk-dominant, as occurs when *w* = 0.5, *P* = 0.5 and *b* = 8. Groups under these conditions evolving in isolation should, all else equal, be just as likely to converge to a TFT-playing equilibrium as to a DIMAS-playing equilibrium. However, since a population of TFT-playing agents would have higher overall payoffs than DIMAS-playing agents, when groups come into competition, TFT-playing groups should out-compete DIMAS-playing groups and one-shot cooperation can spread. Of course, if evolution has access to the entire strategy space we should see many more equilibria than these two. Punishment also can generate multiple equilibria (Boyd and Richerson, 1992).

This process of “equilibrium selection” is generalizable to any case where there is variation between groups and group compete (Boyd and Richerson, 1990) and between-group variation is likely to be greater when behavior is transmitted socially than genetically. This is because when a norm psychology induces humans to adopt their group’s traits, this further reinforces any existing equilibrium and drives down within-group variance relative to between-group variance Henrich (2004); Bell (2010). This models show how reciprocity under uncertainty fits into the larger picture of human evolution: it (with punishment) is a mechanism for generating between-group variation in social norms, individuals adapt to this variation by evolving a predisposition to learn and adopt the prevailing norms of their particular group, this norm psychology further stabilizes between-group variation, and when groups come into competition that group with the more cooperative norms will, all else equal, out-compete groups with less cooperative norms. Thus, as Darwin (1873) writes, “the social and moral qualities would tend slowly to advance and be diffused throughout the world.”

## Acknowledgments

I thank Peter J. Richerson, Richard McElreath, Kyle Joyce, Emily Peffer, Katie Demps, Kari Schroeder, Bruce Winterhalder and members of the UC Davis Cultural Evolution and Human Behavioral Ecology Labs for their comments on earlier drafts of this paper and Andrew Delton and Max Krasnow for their help in recreating their original model. This work was funded, in part, by the post-9/11 G.I. Bill, a Block Grant from the University of California, Davis, and part of this work was conducted at the National Institute for Mathematical and Biological Synthesis, sponsored by the National Science Foundation, the U.S. Department of Homeland Security, and the U.S. Department of Agriculture through NSF Awards EF-0832858 and DBI-1300426, with additional support from The University of Tennessee, Knoxville.

## Appendix A. Frequency of One-Shot Cooperation for All Parameters and Treatments

**Figure A.1:**
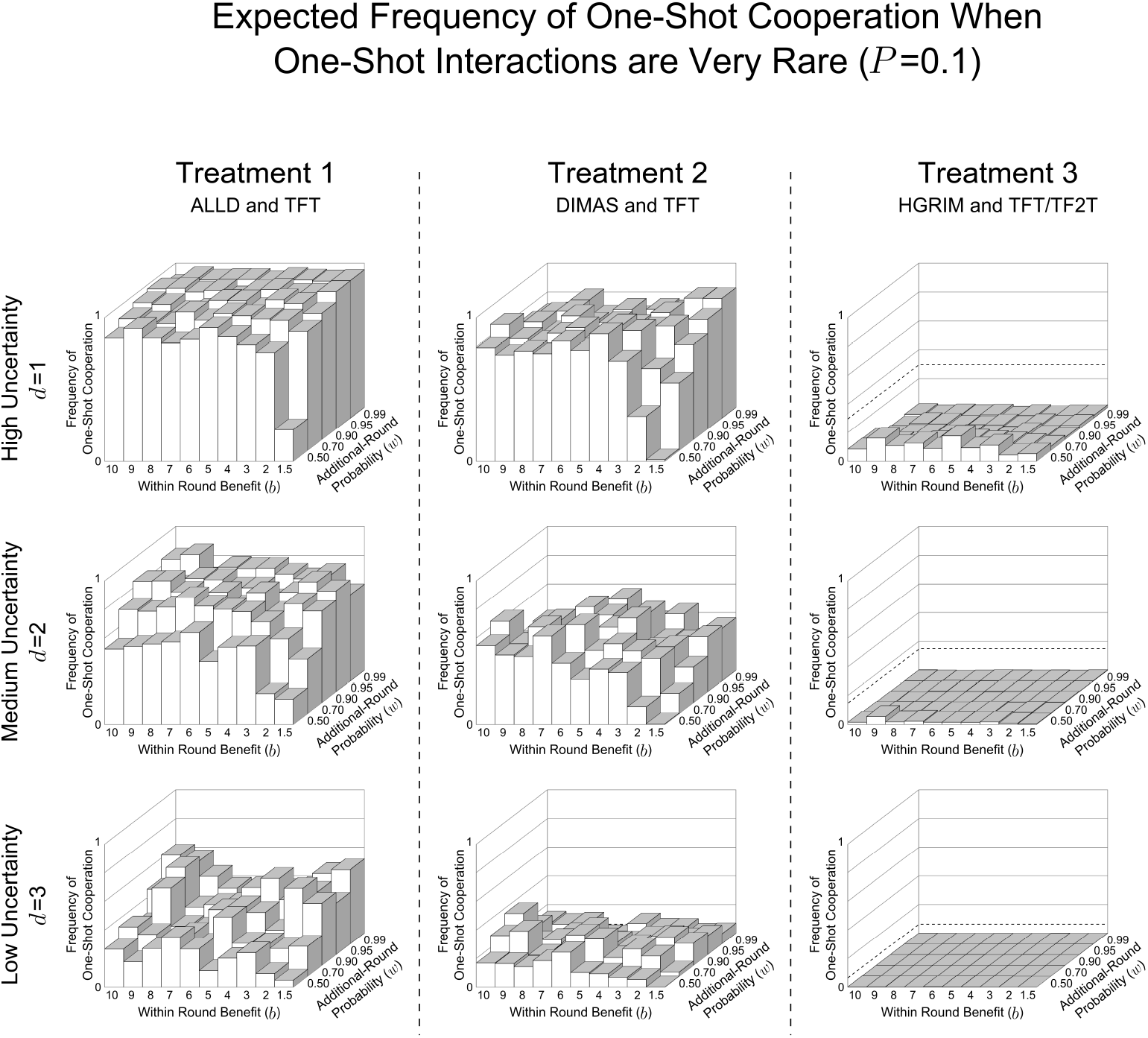
Repentant and forgiving strategies decrease one-shot cooperation well below the DKCT model when one-shot games are very rare (*P* = 0.1) and agents have 10 partners. These show the expected frequency of one-shot cooperation averaged over the last 500 generations of the 10,000 generation simulation for all values of *d*, *b*, and *w*. These are the same parameter combinations reported by DKCT. Treatment 1, where agents, as in DKCT play only TFT or ALLD has the highest frequency of one-shot cooperation. Treatment 2, where ALLD is replaced by a repentant strategy, DIMAS, has less one-shot cooperation. In Treatment 3, where a savvy repentant strategy, HGRIM, competes with TFT and TF2T, one-shot cooperation virtually disappears. This highlights the variability in outcomes in repeated games when one-shot games are very rare.

**Figure A.2:**
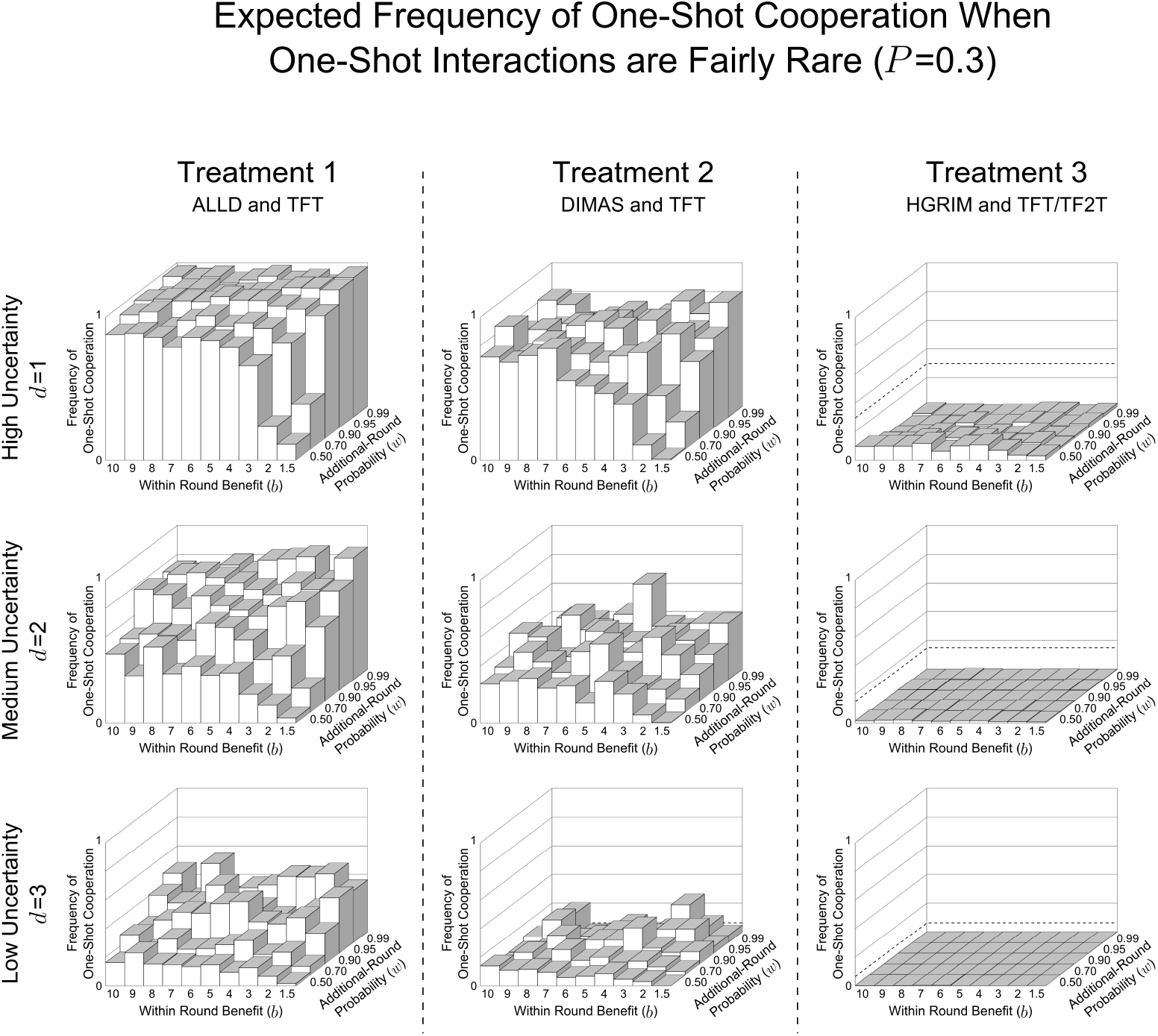
Repentant and forgiving strategies decrease one-shot cooperation well below the DKCT model when one-shot games are fairly rare (*P* = 0.3) and agents have 10 partners. These show the expected frequency of one-shot cooperation averaged over the last 500 generations of the 10,000 generation simulation for all values of *d*, *b*, and *w*. These are the same parameter combinations reported by DKCT. Treatment 1, where agents, as in DKCT play only TFT or ALLD has the highest frequency of one-shot cooperation. Treatment 2, where ALLD is replaced by a repentant strategy, DIMAS, has less one-shot cooperation. In Treatment 3, where a savvy repentant strategy, HGRIM, competes with TFT and TF2T, one-shot cooperation virtually disappears. This highlights the variability in outcomes in repeated games when one-shot games are fairly rare.

**Figure A.3:**
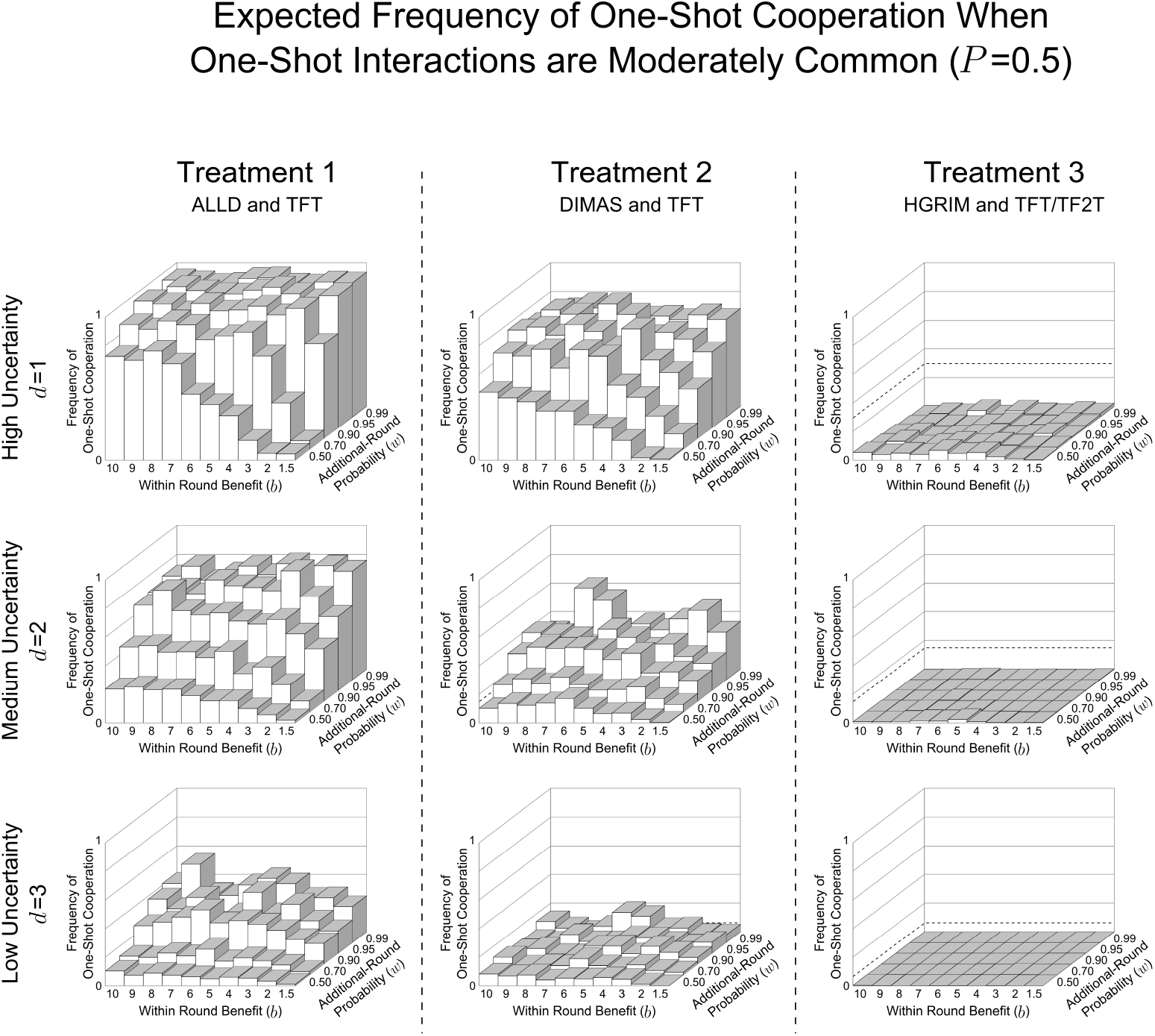
Repentant and forgiving strategies decrease one-shot cooperation well below the DKCT model when one-shot games are moderately rare (*P* = 0.5) and agents have 10 partners. These show the expected frequency of one-shot cooperation averaged over the last 500 generations of the 10,000 generation simulation for all values of *d*, *b*, and *w*. These are the same parameter combinations reported by DKCT. Treatment 1, where agents, as in DKCT play only TFT or ALLD has the highest frequency of one-shot cooperation. Treatment 2, where ALLD is replaced by a repentant strategy, DIMAS, has less one-shot cooperation. In Treatment 3, where a savvy repentant strategy, HGRIM, competes with TFT and TF2T, one-shot cooperation virtually disappears. This highlights the variability in outcomes in repeated games when one-shot games are moderately rare.

**Figure A.4:**
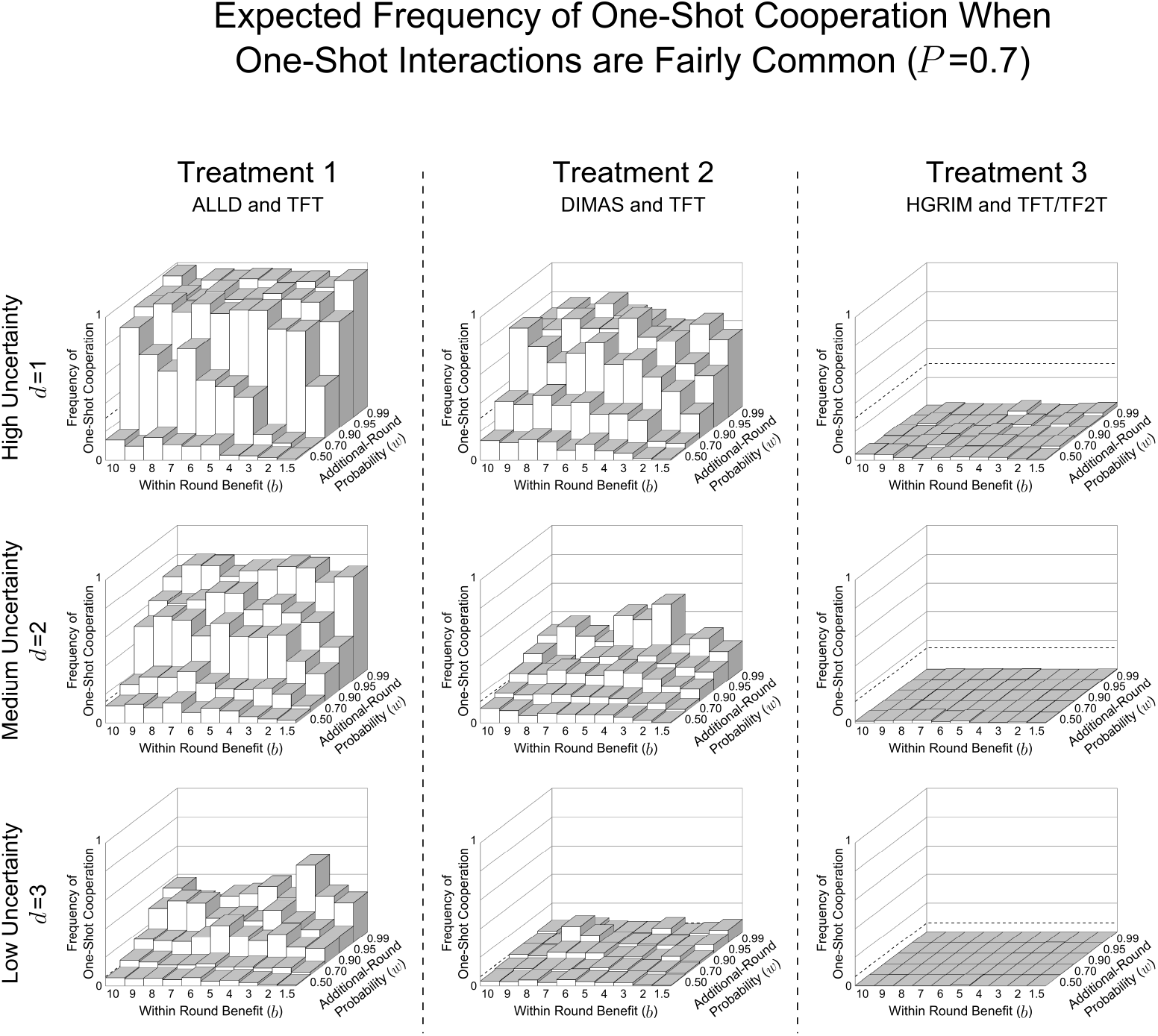
Repentant and forgiving strategies decrease one-shot cooperation well below the DKCT model when one-shot games are fairly common (*P* = 0.7) and agents have 10 partners. These show the expected frequency of one-shot cooperation averaged over the last 500 generations of the 10,000 generation simulation for all values of *d*, *b*, and *w*. These are the same parameter combinations reported by DKCT. Treatment 1, where agents, as in DKCT play only TFT or ALLD has the highest frequency of one-shot cooperation. Treatment 2, where ALLD is replaced by a repentant strategy, DIMAS, has less one-shot cooperation. In Treatment 3, where a savvy repentant strategy, HGRIM, competes with TFT and TF2T, one-shot cooperation virtually disappears. This highlights the variability in outcomes in repeated games when one-shot games are fairly common.

**Figure A.5:**
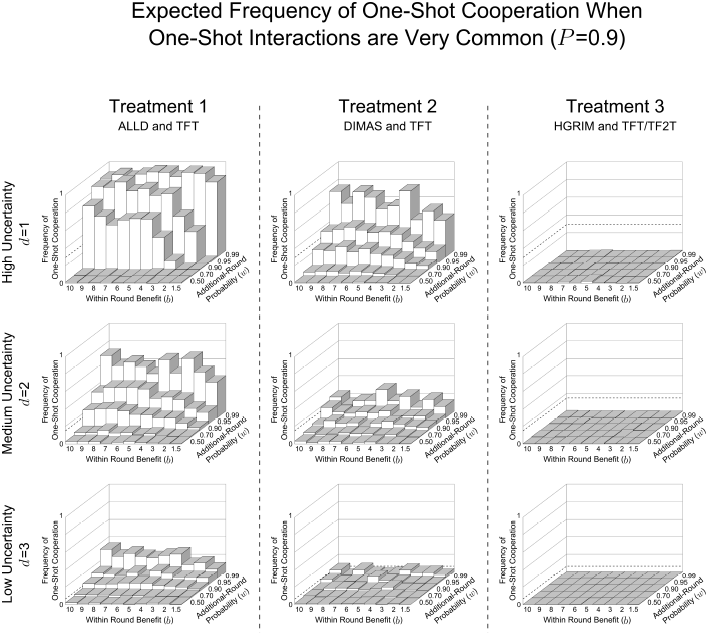
Repentant and forgiving strategies decrease one-shot cooperation well below the DKCT model when one-shot games are very common (*P* = 0.9) and agents have 10 partners. These show the expected frequency of one-shot cooperation averaged over the last 500 generations of the 10,000 generation simulation for all values of *d*, *b*, and *w*. These are the same parameter combinations reported by DKCT. Treatment 1, where agents, as in DKCT play only TFT or ALLD has the highest frequency of one-shot cooperation. Treatment 2, where ALLD is replaced by a repentant strategy, DIMAS, has less one-shot cooperation. In Treatment 3, where a savvy repentant strategy, HGRIM, competes with TFT and TF2T, one-shot cooperation virtually disappears. This highlights the variability in outcomes in repeated games when one-shot games are very common.

**Figure A.6:**
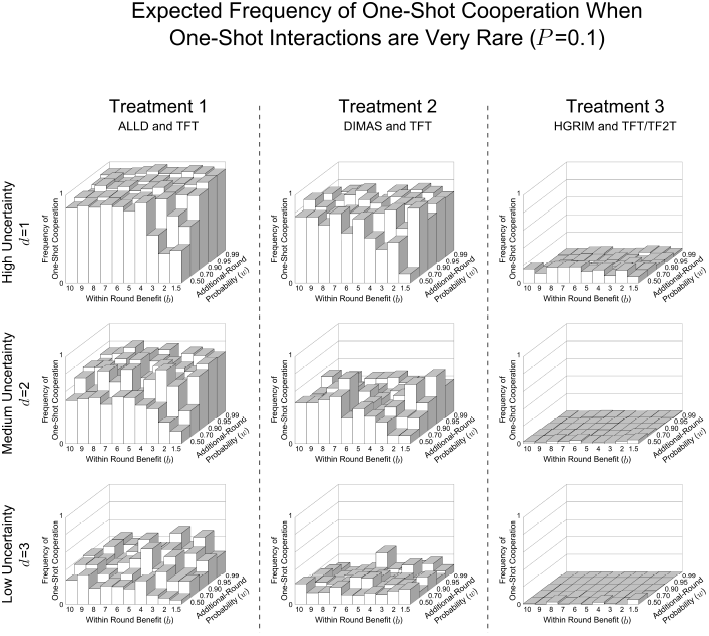
Repentant and forgiving strategies decrease one-shot cooperation well below the DKCT model when one-shot games are very rare (*P* = 0.1) and agents have 10 partners. These show the expected frequency of one-shot cooperation averaged over the last 500 generations of the 10,000 generation simulation for all values of *d*, *b*, and *w*. These are the same parameter combinations reported by DKCT. Treatment 1, where agents, as in DKCT play only TFT or ALLD has the highest frequency of one-shot cooperation. Treatment 2, where ALLD is replaced by a repentant strategy, DIMAS, has less one-shot cooperation. In Treatment 3, where a savvy repentant strategy, HGRIM, competes with TFT and TF2T, one-shot cooperation virtually disappears. This highlights the variability in outcomes in repeated games when one-shot games are very rare.

**Figure A.7:**
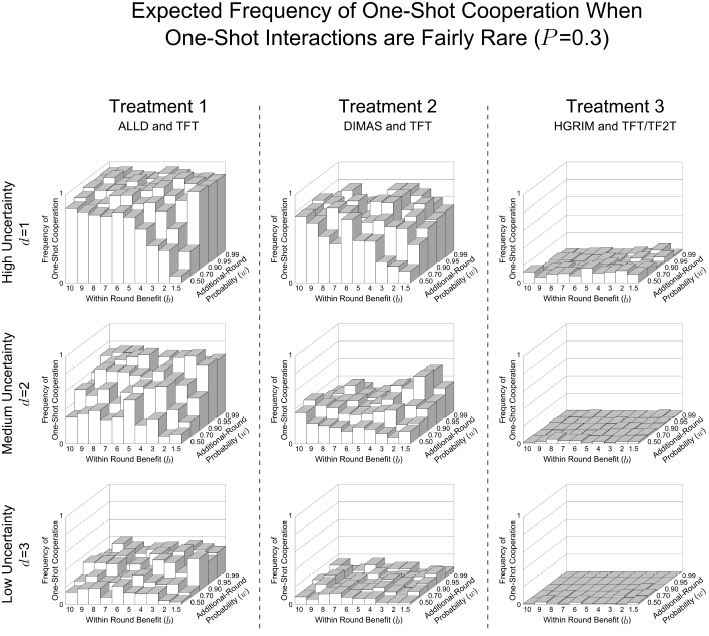
Repentant and forgiving strategies decrease one-shot cooperation well below the DKCT model when one-shot games are fairly rare (*P* = 0.3) and agents have one partner. These show the expected frequency of one-shot cooperation averaged over the last 500 generations of the 10,000 generation simulation for all values of *d*, *b*, and *w*. These are the same parameter combinations reported by DKCT. Treatment 1, where agents, as in DKCT play only TFT or ALLD has the highest frequency of one-shot cooperation. Treatment 2, where ALLD is replaced by a repentant strategy, DIMAS, has less one-shot cooperation. In Treatment 3, where a savvy repentant strategy, HGRIM, competes with TFT and TF2T, one-shot cooperation virtually disappears. This highlights the variability in outcomes in repeated games when one-shot games are fairly rare.

**Figure A.8:**
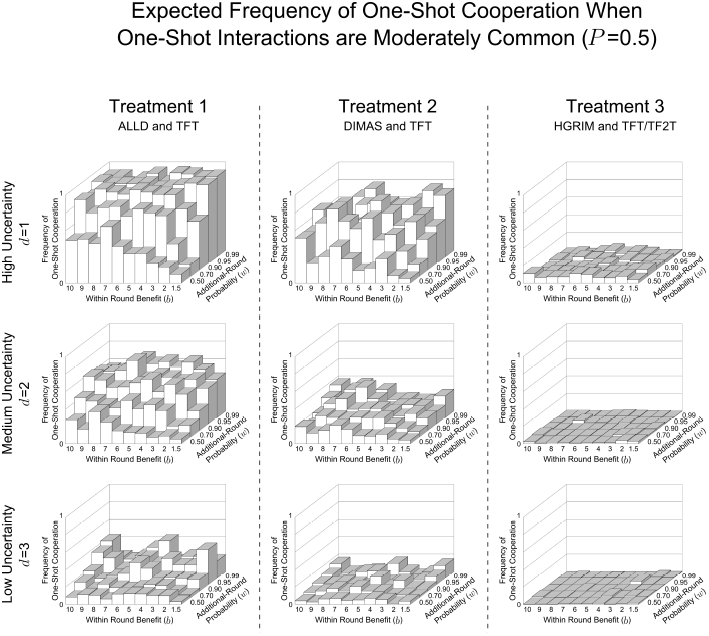
Repentant and forgiving strategies decrease one-shot cooperation well below the DKCT model when one-shot games are moderately rare (*P* = 0.5) and agents have one partner. These show the expected frequency of one-shot cooperation averaged over the last 500 generations of the 10,000 generation simulation for all values of *d*, *b*, and *w*. These are the same parameter combinations reported by DKCT. Treatment 1, where agents, as in DKCT play only TFT or ALLD has the highest frequency of one-shot cooperation. Treatment 2, where ALLD is replaced by a repentant strategy, DIMAS, has less one-shot cooperation. In Treatment 3, where a savvy repentant strategy, HGRIM, competes with TFT and TF2T, one-shot cooperation virtually disappears. This highlights the variability in outcomes in repeated games when one-shot games are moderately rare.

**Figure A.9:**
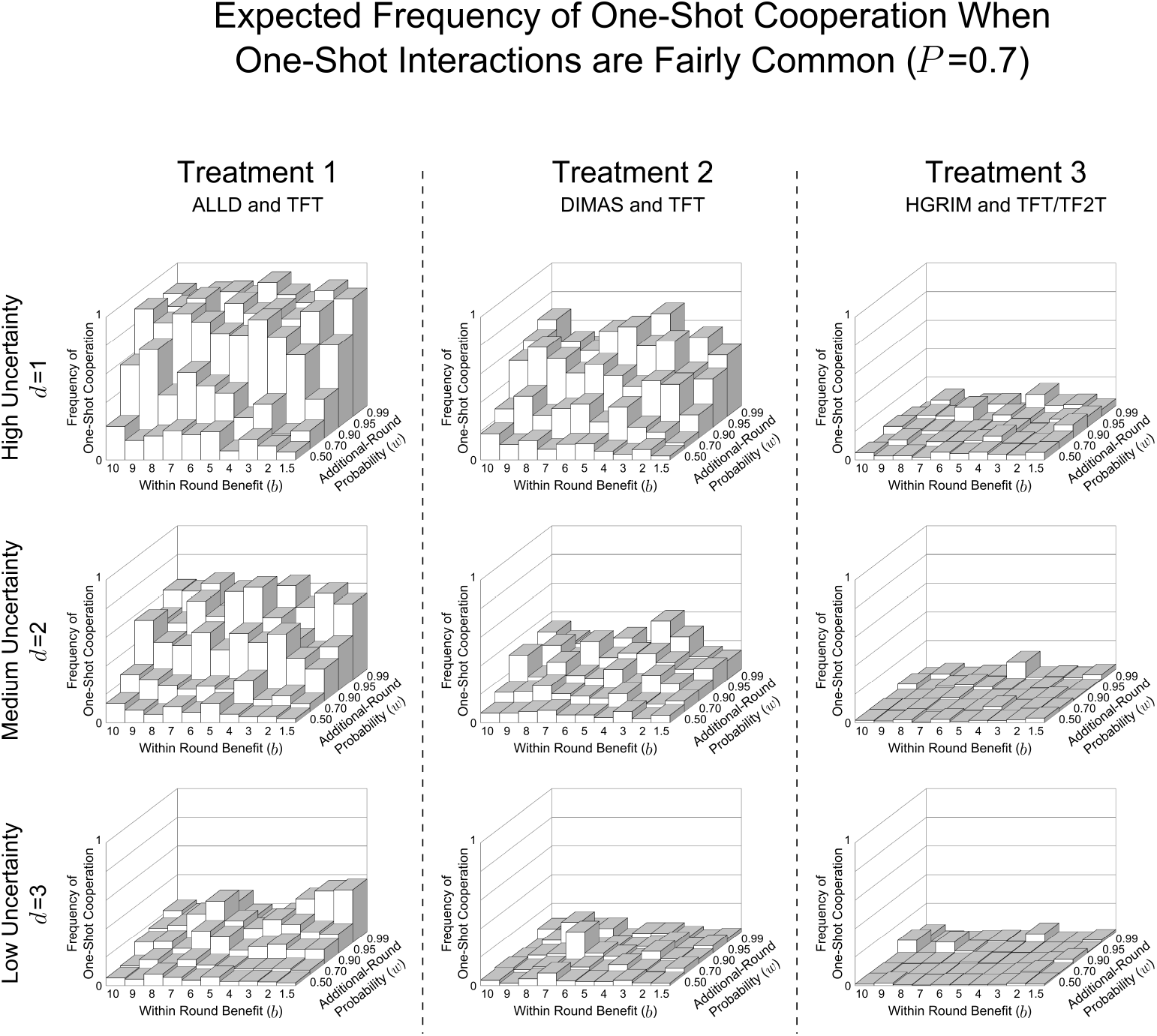
Repentant and forgiving strategies decrease one-shot cooperation well below the DKCT model when one-shot games are fairly common (*P* = 0.7) and agents have one partner. These show the expected frequency of one-shot cooperation averaged over the last 500 generations of the 10,000 generation simulation for all values of *d*, *b*, and *w*. These are the same parameter combinations reported by DKCT. Treatment 1, where agents, as in DKCT play only TFT or ALLD has the highest frequency of one-shot cooperation. Treatment 2, where ALLD is replaced by a repentant strategy, DIMAS, has less one-shot cooperation. In Treatment 3, where a savvy repentant strategy, HGRIM, competes with TFT and TF2T, one-shot cooperation virtually disappears. This highlights the variability in outcomes in repeated games when one-shot games are fairly common.

**Figure A.10:**
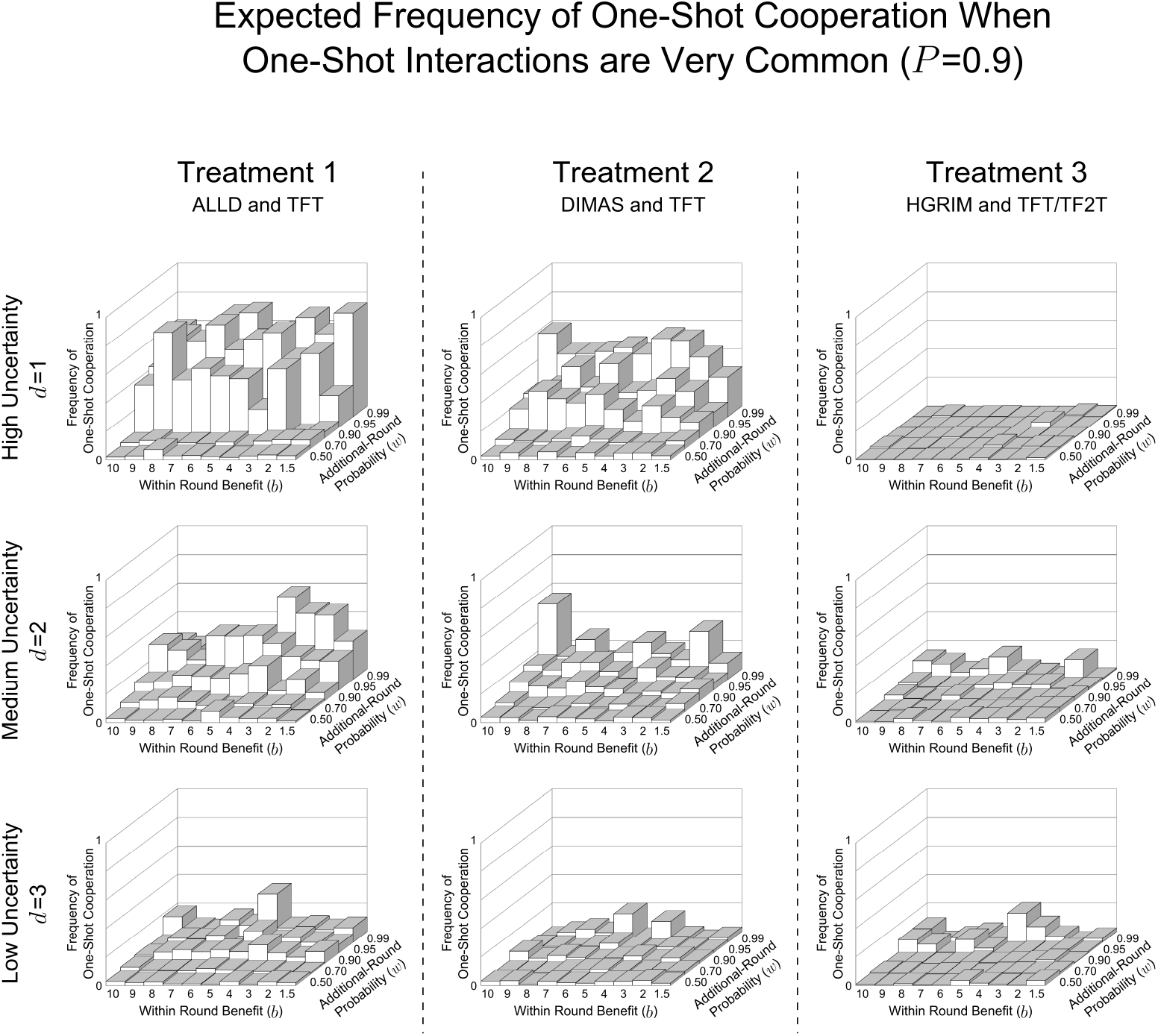
Repentant and forgiving strategies decrease one-shot cooperation well below the DKCT model when one-shot games are very common (*P* = 0.9) and agents have one partner. These show the expected frequency of one-shot cooperation averaged over the last 500 generations of the 10,000 generation simulation for all values of *d*, *b*, and *w*. These are the same parameter combinations reported by DKCT. Treatment 1, where agents, as in DKCT play only TFT or ALLD has the highest frequency of one-shot cooperation. Treatment 2, where ALLD is replaced by a repentant strategy, DIMAS, has less one-shot cooperation. In Treatment 3, where a savvy repentant strategy, HGRIM, competes with TFT and TF2T, one-shot cooperation virtually disappears. This highlights the variability in outcomes in repeated games when one-shot games are very common.

## Appendix B. Complete Uncertainty Calculations

Under complete uncertainty, DKCT’s model reduces to a standard repeated Prisoner’s Dilemma where the probability of transitioning from the first to second round is lower, (1 − *P)w*, than the transition probability for every other round, *w*. Figure B.1 gives the expected payoffs for the strategy combinations in all three treatments under complete uncertainty. In this Appendix I show the calculation used to derive the stability and risk-dominance conditions in Table 3.

**Figure B.1:**
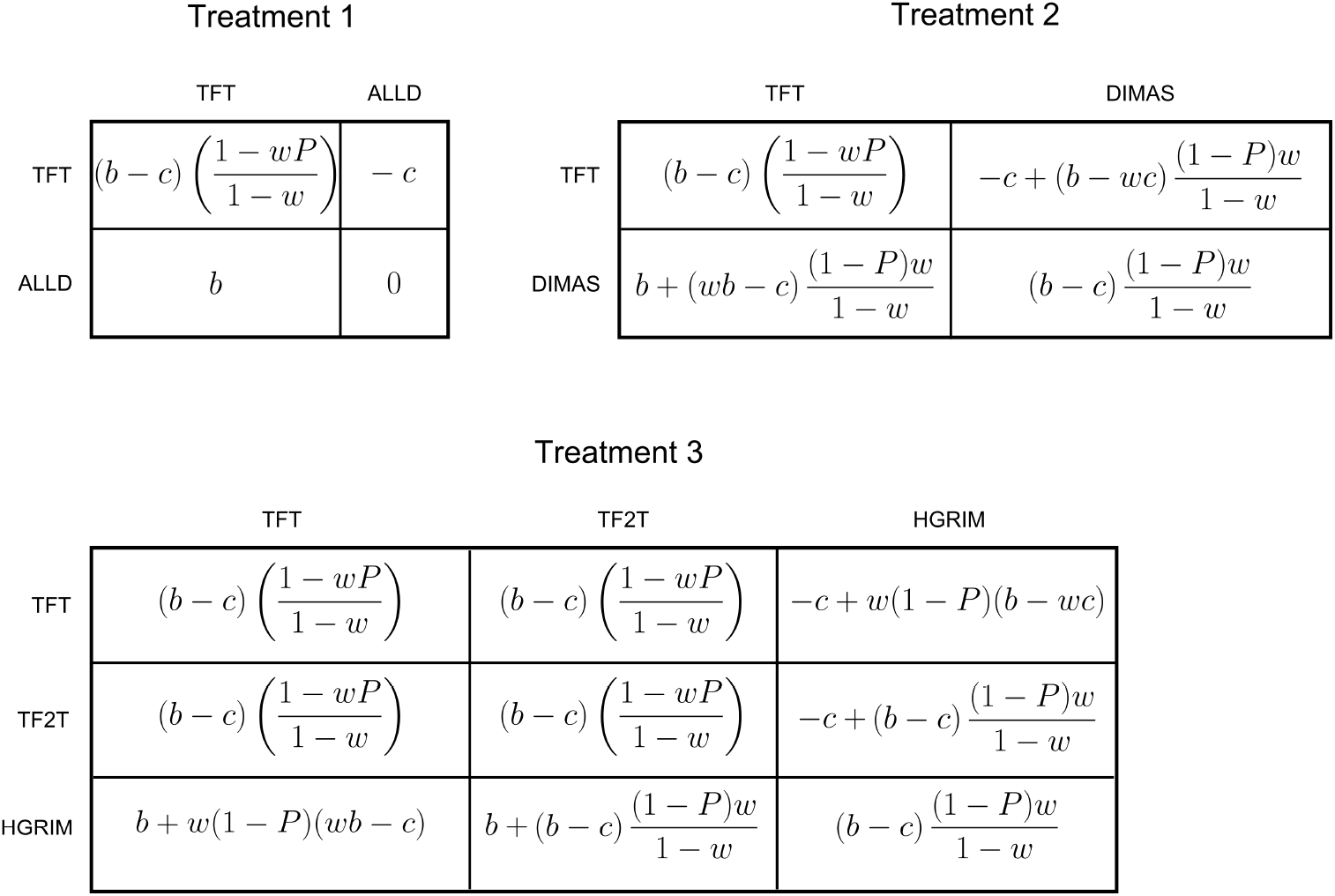
Expected payoffs for strategies in the three treatments when agents are completely uncertain whether a game is one-shot or repeated. This is the best-case scenario for the evolution of one-shot altruistic cooperation. *b* is the benefits to cooperation, *c* is the cost of cooperating, *P* is the probability that an interaction is one-shot, and *w* is the probability that, after each round in a repeated game, there is another round.

### Appendix B.1. When is TFT stable against ALLD?

TFT is stable against ALLD when the payoff to TFT given TFT is greater than the payoff to ALLD given TFT:

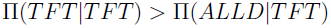

Substituting from Figure B.1 and simplifying yields the condition where TFT, and thus one-shot cooperation is stable:

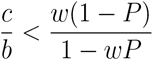

### Appendix B.2. When is TFT stable against DIMAS?

TFT is stable against DIMAS when the payoff to TFT given TFT is greater than the payoff to DIMAS given TFT:

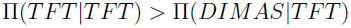

Substituting from Figure B.1 and simplifying yields the condition where TFT, and thus one-shot cooperation is stable:

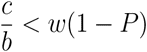

### Appendix B.3. When is TFT stable against direct invasion by HGRIM?

TFT is stable against HGRIM when the payoff to TFT given TFT is greater than the payoff to HGRIM given TFT:

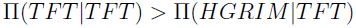

Substituting and Simplifying:

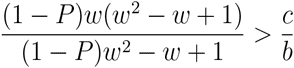

### Appendix B.4. When is TFT stable against indirect invasion by HGRIM via TF2T?

One of two conditions must be met for TFT to be stable against indirect invasion by HGRIM via TF2T. First the payoff to TFT given HGRIM could be greater than the payoff to TF2T given HGRIM:

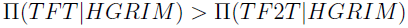

Substituting from Figure B.1 and simplifying yields the condition where TFT, and thus one-shot cooperation is stable:

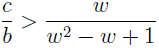

Second the payoff to TF2T given HGRIM may be greater than the payoff to TF2T given TF2T. By inspection of Figure B.1 this can never be true because their payoffs are equivalent in all rounds except the first where TF2T pays a cost of *-c*.

### Appendix B.5. When is TFT stable against both direct and indirect invasion by HGRIM?

From Appendix B.3 and Appendix B.4, the condition where TFT is stable against both direct and indirect invasion is:

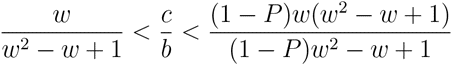

It is easy to show that 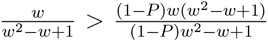 over all possible values of *P*, *w*, and 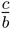, therefore TFT is never stable against co-invasion by TFT and HGRIM.

## Appendix C. Effects of Increasing Partners from One to Ten

Figure C.1A shows the typical distribution of the average number rounds per partner for each agent in a generation when *P* = 0.9 and *w* = 0.99 when agents have only partners. With only one partner, 92% of the agents play only one round in their lifetime. One dyad plays well over 500 rounds, more rounds than the rest of the agents combined. Over the course of 10,000 generations, a dyad of agents playing poorly performing strategies will sometimes be assigned one of these outliers and their strategies can go to near fixation.

In Figure C.1B agents play the same game with ten partners. The distribution of the number of rounds per partner averages out agents who play a game with a disproportionally high number of rounds with one partner are likely to play games of much fewer rounds with another. Here 58% of agents average two or more rounds per partner and the greatest outlier is much closer to the mean number of rounds. (As the number of partners increases, the average number of rounds per partner for each agent should approach the average number of rounds of the distribution, 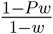). Playing with multiple partners limits the effect of stochastic shocks while still maintaining the logical structure of the game itself.

**Figure C.1:**
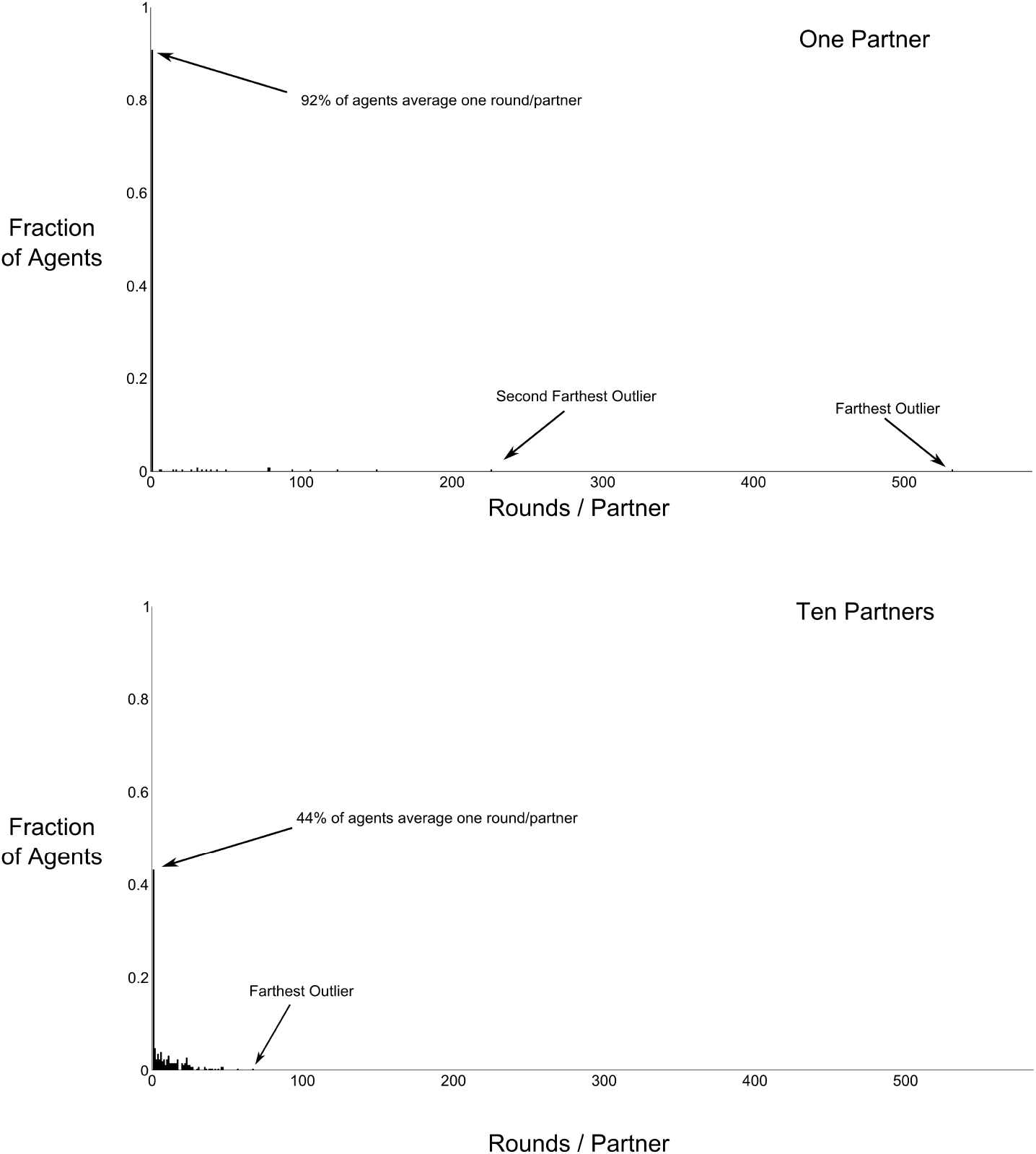
Increasing the number of partners per player decreases the amount of payoff variance due to number of rounds played. A shows a typical distribution of rounds of play per partner for each of 500 players for conditions *P* = 0.1 and *w* = 0.99. B shows a typical distribution of rounds of play per partner under the same conditions, if each agent has 10 partners in their lifetime.

## Appendix D. Finite State Machine Representations of Strategies

Sometimes verbal descriptions of strategies can be ambiguous. Strategies from all three treatments are represented in Figure D.1 as Finite State Automata. Initial plays of the strategy are represented by the state in the double circle (nasty strategies start with defect and nice strategies start with cooperate). Transition rules to the next state are represented by arrows. For example, DIMAS starts with Defect, transitions to Cooperate, and stays at Cooperate until its opponent defects. After defection by an opponent, DIMAS

**Figure D.1:**
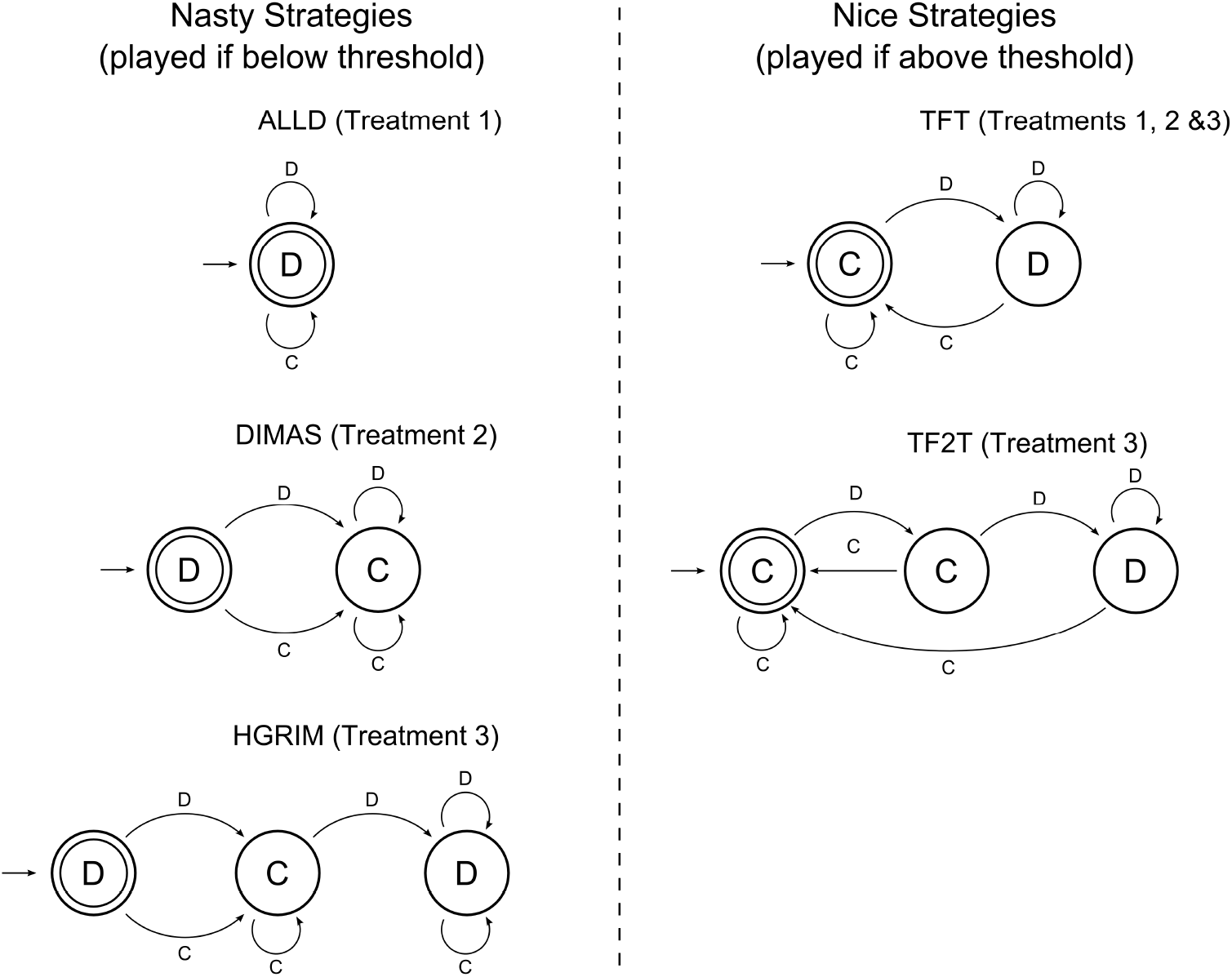
The strategies included in all three treatments represented as Moore Machines, a class of Finite State Automata. Then it defects forever. The number of states in the minimal FSA is a measure of the memory of a strategy. ALLD is a memory-one strategy. TFT and DIMAS are memory-two strategies. TF2T and HGRIM are memory-three strategies.

## Footnotes

1 Though it has also been called the “Savanna Principle” (Kanazawa, 2004), the “big mistake hypothesis” (Richerson and Boyd, 2005), the “misapprehension hypothesis” (Hagen and Hammerstein, 2006), the “evolutionary legacy hypothesis” (Burnham and Johnson, 2005) and “social exchange theory” (Krasnow et al., 2012).

2 DCKT also report similar results from a version of the model with the same game structure and strategy space, but where agents have more complicated cognitive architecture. Since the logic of payoff-based selection applies to any cognitive architecture that allows for a sufficient range of possible strategies, for tractability I focus on the simpler model.

3 It is called a “folk theorem” because, while understood by game theorists by the 1950s and 1960s, it is unknown who derived it.

4 The biblical Dimas’s repentance, afterall, was only successful because of Jesus’s forgiveness.

5 TF2T, of course, did not win the second tournament, which further highlights that a strategy’s success hinges on the mix of other strategies in the population.

6 Though in practice, the long run might be *very* long.

